# Requirement of sequential hydrolysis by CD73 and ALP for uptake of vitamin B_2_ into cells

**DOI:** 10.1101/2022.07.20.500786

**Authors:** Natsuki Shichinohe, Daisuke Kobayashi, Ayaka Izumi, Kazuya Hatanaka, Rio Fujita, Taroh Kinoshita, Norimitsu Inoue, Naoya Hamaue, Keiji Wada, Yoshiko Murakami

**Affiliations:** Department of Food and Chemical Toxicology, School of Pharmaceutical Sciences, Health Sciences University of Hokkaido, Ishikari-Tobetsu, Hokkaido 061-0293, Japan; Laboratory of Immunoglycobiology, Research Institute for Microbial Diseases, Osaka University, Suita, Osaka 565-0871, Japan; WPI Immunology Frontier Research Center, Osaka University, Suita, Osaka 565-0871, Japan; Center for Infectious Disease Education and Research, Osaka University, Suita, Osaka 565-0871, Japan; Department of Molecular Genetics, Wakayama Medical University, Wakayama, Wakayama 641-8509, Japan

**Keywords:** Alkaline phosphatase, CD73, FAD, GPI anchored protein, Mitochondrial function

## Abstract

Extracellular hydrolysis of flavin adenine dinucleotide (FAD) and flavin mononucleotide (FMN) to riboflavin is thought to be important for cellular uptake of vitamin B_2_ because FAD and FMN are hydrophilic and do not pass the plasma membrane. However, it is not clear whether FAD and FMN are hydrolyzed by cell surface enzymes for vitamin B_2_ uptake. Here, we show that in human cells, FAD, a major form of vitamin B_2_ in plasma, is hydrolyzed by CD73 (also called ecto-5′ nucleotidase) to FMN, then FMN is hydrolyzed by alkaline phosphatase to riboflavin, which is efficiently imported into cells. This process is impaired on the surface of glycosylphosphatidylinositol (GPI)-deficient cells due to lack of these GPI-anchored enzymes. During culture of GPI-deficient cells with FAD or FMN, hydrolysis of these forms of vitamin B_2_, intracellular levels of vitamin B_2_, vitamin B_2_-dependent pyridoxal 5′-phosphate formation, and mitochondrial functions were significantly decreased compared with those in GPI-restored cells. These results suggest that inefficient uptake of vitamin B_2_ might account for mitochondrial dysfunction seen in some cases of inherited GPI deficiency.

## Introduction

The water-soluble vitamins are essential to mammals and play roles in various metabolic reactions by working as coenzymes. The major forms of vitamins B_1_, B_2_, and B_6_ in blood are phosphorylated or nucleotide forms (Rindi *et al*, 1981; Hustad *et al*, 1999; Vasilaki *et al*, 2010; Talwar *et al*, 2003; Ueland *et al*, 2015; Akiyama *et al*, 2017) and they need to be hydrolyzed for uptake by cells. Alkaline phosphatase (ALP), a glycosylphosphatidylinositol (GPI)-anchored protein (GPI-AP), dephosphorylates thiamine pyrophosphate and pyridoxal 5′-phosphate (PLP) to thiamine and pyridoxal (PL), respectively (Luong & Nguyen, 2005; Millán & Whyte, 2016). However, it is uncertain whether the active forms of vitamin B_2_, flavin adenine dinucleotide (FAD, Fig 1A) and flavin mononucleotide (FMN, Fig 1A), are hydrolyzed by the cell surface enzyme for their uptake. Moreover, enzymes which hydrolyze them, are not known (Barile *et al*, 2016).

**Figure 1.**
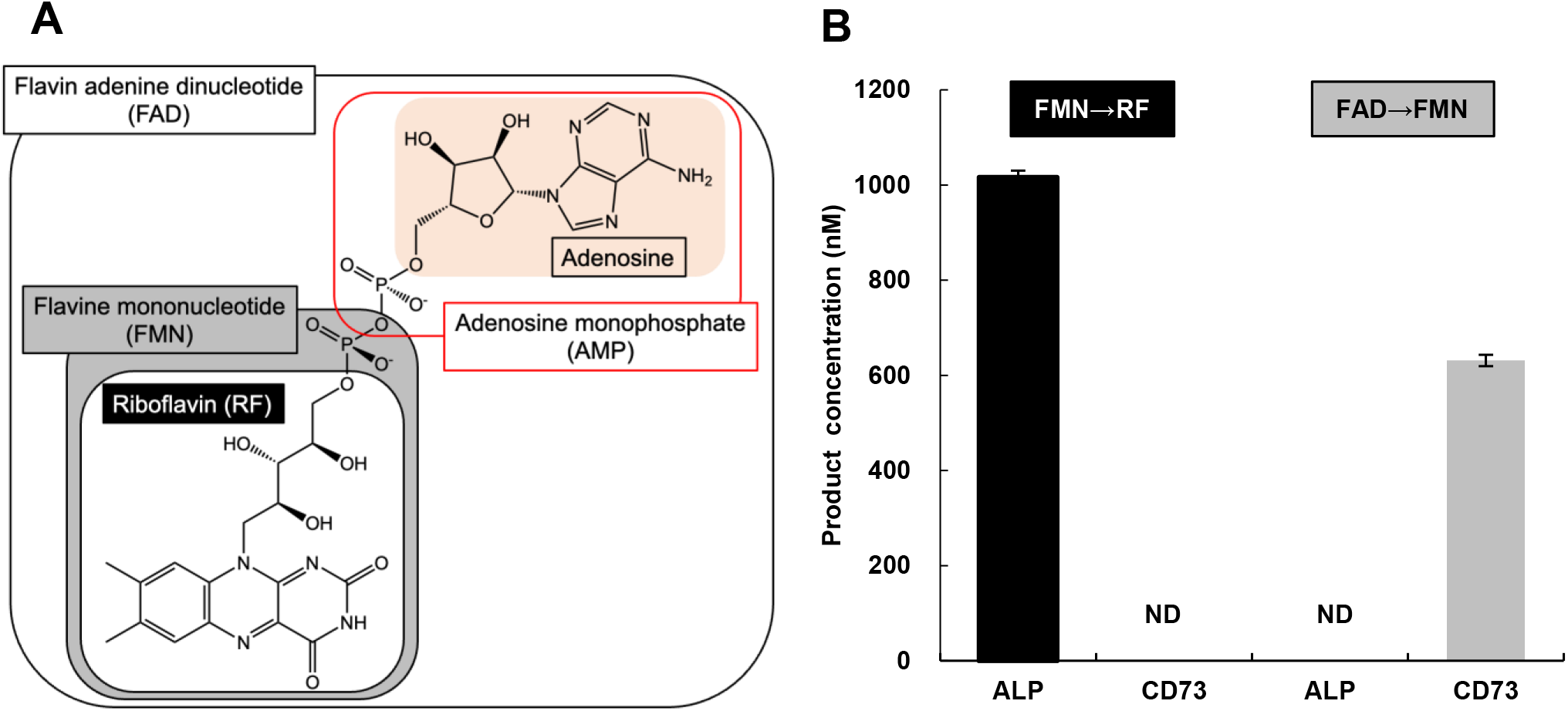
Structures of vitamin B_2_ analogues and hydrolysis of flavin mononucleotide (FMN) and flavin adenine dinucleotide (FAD) hydrolysis. A. Structures of FAD and metabolites of FAD B. FMN and FAD hydrolysis by human bone alkaline phosphatase (ALP) and recombinant human CD73 Riboflavin (RF) and FMN concentrations were measured 15 min after incubation of 10 μM FMN or FAD, respectively, with ALP (10 μg/mL) or CD73 (10 ng/mL). Data represent means ± S.D. (*n* = 3). N.D.: Not detectable.

In food, vitamin B_2_ is mainly in the form of FAD or FMN. Because they are hydrophilic, they are not imported directly into the intestinal epithelial cells; instead, they must be converted to riboflavin (RF, Fig 1A) on cell surface for uptake (Akiyama *et al*, 1982; Daniel *et al*, 1983). RF uptake into cells is mediated by three kinds of RF transporters (SLC52A1, 2, 3, Barile *et al,* 2016). FMN and FAD are then regenerated from RF in the cytoplasm by riboflavin kinase and FAD synthetase (Said & Arianas, 1991; Barile *et al*, 2016). These active forms of vitamin B_2_ work as cofactors of various enzymes involved in redox reactions in many metabolic pathways, such as the tricarboxylic acid cycle, vitamin B_6_ metabolism and mitochondrial electron transport chain. Therefore, vitamin B_2_ deficiency causes mitochondrial dysfunction as well as various metabolic disorders (Barile *et al*, 2016).

One candidate enzyme involved in the hydrolysis of FMN and FAD is ALP. Daniel et al reported that FMN and FAD were hydrolyzed by ALP purified from the brush-border membrane of rat jejunum (Daniel *et al*, 1983). ALP hydrolyzes compounds with a phosphate moiety, such as pyrophosphate, phosphoethanolamine, and PLP (Millán & Whyte, 2016). There are four main isoforms of ALP in humans, — tissue nonspecific ALP (TNSALP), intestinal ALP, germ cell ALP, and placental ALP (PLAP)—all of which are GPI-APs (Zimmermann *et al*, 2012). Among them, TNSALP is ubiquitously expressed and is the major isoform expressed in liver, bone, kidney, blood, and brain (Millán, 2006).

There are several reports suggesting that two enzymes are involved in hydrolysis of FMN and FAD. Akiyama *et al*. purified two independent enzymes, FAD pyrophosphatase and FMN phosphatase, from rat intestinal brush-border (Akiyama *et al*, 1982). Okuda showed that the inhibitory effect of pyrophosphate was greater against FMN hydrolysis than FAD hydrolysis in dog intestinal mucosa, and suggested that the small intestine contains at least two kinds of phosphatase: nucleotide pyrophosphatase with low-affinity for pyrophosphate and phosphomonoesterase with high-affinity for pyrophosphate (Okuda, 1958a). Okuda also reported that gastric juice hydrolyzes only FAD, and bile and pancreatic juice hydrolyze only FMN (Okuda, 1958b). Lee *et al*. showed that an enzyme purified from placental trophoblastic microvilli possessed FAD pyrophosphatase activity and the enzyme did not hydrolyze FMN (Lee & Ford, 1988). These reports suggest a two-step hydrolysis of FAD (i.e., FAD to FMN, then FMN to RF).

Here, we show that, in human cells, FAD is hydrolyzed in two steps: FAD to FMN by CD73, and FMN to RF by ALP. CD73, also known as ecto-5′ nucleotidase, is a GPI-AP encoded by the *NT5E* gene and catalyzes conversion of AMP to adenosine (Picher *et al*, 2003; Zimmermann *et al*, 2012). To determine the hydrolytic activity of human bone ALP and recombinant human CD73 toward vitamin B_2_ analogues, RF formation from FMN and FMN formation from FAD were measured. The contributions of these GPI-APs to vitamin B_2_ uptake, vitamin B_2_-dependent vitamin B_6_ metabolism, and mitochondrial function were determined using GPI-deficient cells.

Because CD73 and ALP are both GPI-APs (Zimmermann *et al*, 2012; Misumi *et al*, 1990), GPI-deficient cells contained a decreased amount of intracellular vitamin B_2_ when cultured with FAD as the vitamin B_2_ source, leading to changes in vitamin B_6_ metabolism and mitochondrial dysfunction. These lines of evidence suggest that the mitochondrial dysfunction seen in some severe cases of inherited GPI deficiency (IGD) might be caused by vitamin B_2_ deficiency, which would be prevented by administration of RF.

## Results

### RF formation from FMN by ALP and FMN formation from FAD by CD73

To analyze the two-step hydrolysis of FAD, RF and FMN concentrations were measured *in vitro* by high-performance liquid chromatography (HPLC) after incubation of 10 μM FMN and FAD, respectively, with human bone ALP and CD73 containing a C-terminal His tag in solution for 15 min. An increase in the RF concentration was detected after incubation of FMN with ALP but not with CD73. In contrast, the FMN concentration was increased after incubation of FAD with CD73 but not with ALP (Fig 1B).

ALP-dependent RF formation from FMN is shown in Fig 2A–C, including a time course and the dependence on the concentrations of ALP and FMN, respectively. RF production increased linearly for 10 min (Fig 2A) in an ALP concentration-dependent manner (Fig 2B). RF production from FMN by ALP was substrate-saturable (Fig 2C), the *Km* and *Vmax* values being 0.309 ± 0.051 μM and 7.47 ± 0.25 nmol/min/mg protein, respectively (Table 1). These results demonstrate that purified bone ALP, that is TNSALP, hydrolyzes FMN, producing RF.

**Figure 2.**
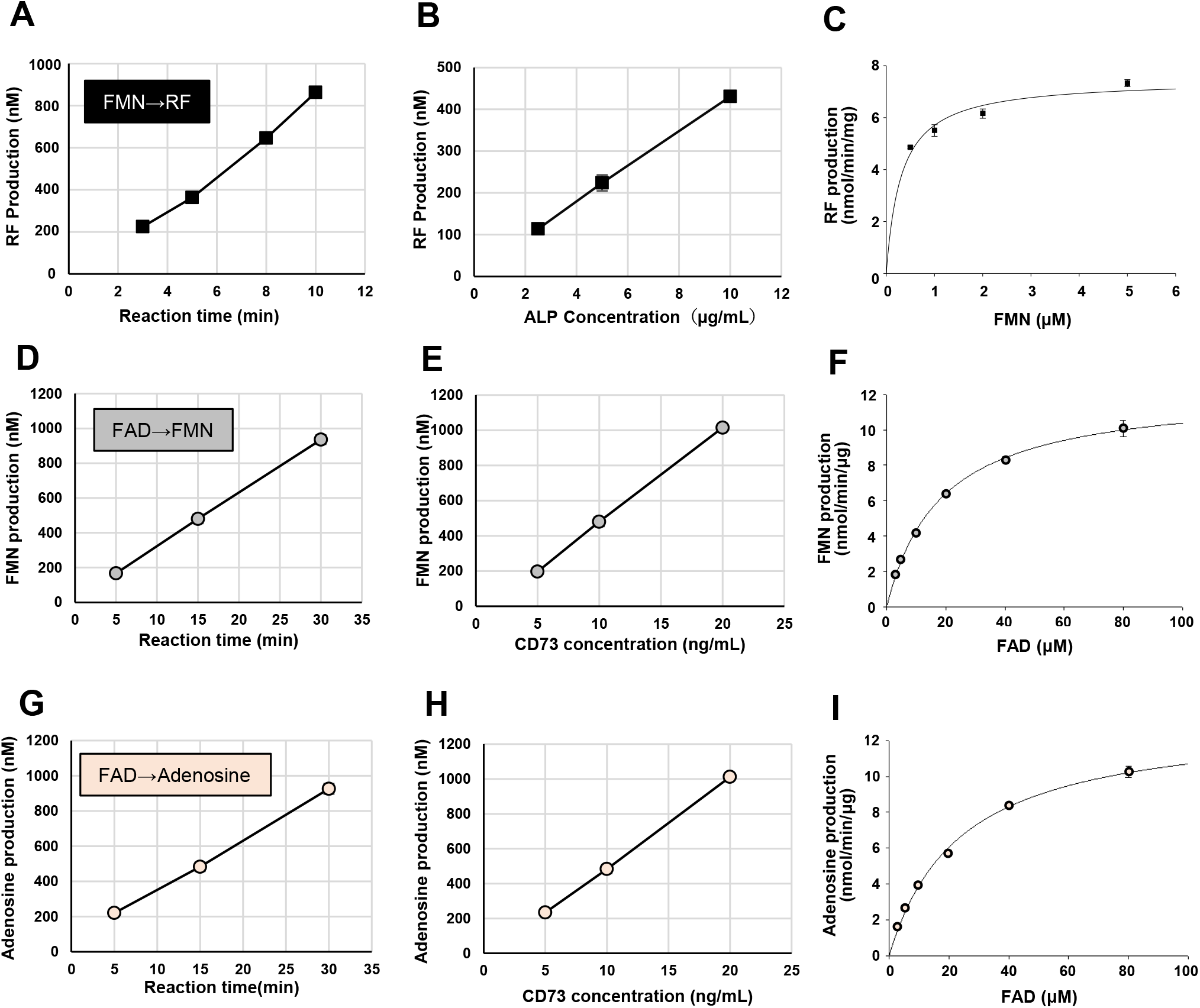
FMN hydrolysis by ALP (A–C) and FAD hydrolysis by CD73 (D–I) A. Time course of RF production from FMN by ALP. The RF concentration was measured after incubation of 10 μM FMN with ALP (10 μg/mL). Data represent means ± S.D. (*n* = 3). B. ALP concentration-dependence of RF production from FMN. The RF concentration was measured 5 min after incubation of 10 μM FMN with ALP (2.5–10 μg/mL). Data represent means ± S.D. (*n* = 3). C. Michaelis-Menten plot of RF production from FMN by ALP. The RF concentration was measured 5 min after incubation of FMN (0.5–5 μM) with ALP (5 μg/mL). Data represent means ± S.D. (*n* = 3). D, G. Time course of FMN production (D) or adenosine production (G) from FAD by CD73. The FMN or adenosine concentration was measured after incubation of 10 μM FAD with CD73 (10 ng/mL). Data represent means ± S.D. (*n* = 3). E, H. CD73 concentration-dependence of FMN production (E) or adenosine production (H) from FAD by CD73. The FMN or adenosine concentration was measured 15 min after incubation of 10 μM FAD with CD73 (5–20 ng/mL). Data represent means ± S.D. (*n* = 3). F, I. Michaelis-Menten plot of FMN production (F) or adenosine production (I) from FAD by CD73. The FMN or adenosine concentration was measured 15 min after incubation of 10 μM FAD with CD73 (10 ng/mL). Data represent means ± S.D. (*n* = 3). When the S.D. is smaller than the symbols, it is not shown.

**Table 1.** Kinetic parameters for hydrolysis reactions catalyzed by ALP and CD73 Data represent means ± standard errors.

a) ALP
| | *K<sub>m</sub>*<br>(μM) | *V<sub>max</sub>*<br>(nmol/min/mg protein) |
| --- | --- | --- |
| FMN → RF | 0.309 ± 0.051 | 7.47 ± 0.25 |
| AMP → Adenosine | 0.538 ± 0.047 | 13.8 ± 0.4 |

b) CD73
| | *K<sub>m</sub>*<br>(μM) | *V<sub>max</sub>*<br>(nmol/min/μg protein) |
| --- | --- | --- |
| FAD → FMN | 18.4 ± 0.3 | 12.3 ± 0.3 |
| FAD → Adenosine | 23.7 ± 1.5 | 13.2 ± 0.3 |
| AMP → Adenosine | 3.94 ± 0.51 | 188 ± 7 |

Recombinant CD73-dependent FMN formation from FAD is shown in Fig 2D–F. Because FAD consists of RF, diphosphate and adenosine moieties, AMP (adenosine + phosphate) or adenosine could be produced by FAD hydrolysis (Fig 1A). To determine the products of FAD hydrolysis by CD73, we simultaneously determined FMN, AMP, and adenosine by HPLC after incubation of FAD with CD73. Increased concentrations of FMN and adenosine were detected, but no increase in AMP was observed. The concentrations of FMN and adenosine increased linearly for 30 min (Fig 2D, G) in a CD73 concentration-dependent manner (Fig 2E, H). FMN and adenosine production by CD73 was dependent on the concentration of the substrate, FAD, and was saturable (Fig 2F, I). CD73 is known to hydrolyze AMP to adenosine; in addition, here, we show evidence that recombinant CD73 also hydrolyzes FAD to produce FMN and adenosine.

### Kinetic parameters of ALP and CD73 activities

We determined kinetic parameters of the activities of ALP and CD73 (Table 1). Because AMP is a substrate for both CD73 and ALP (Picher *et al*, 2003; Zimmermann *et al*, 2012), the results for kinetic analysis of FMN and FAD hydrolysis are compared with those for AMP hydrolysis in Table 1. The *Km* value for FAD hydrolysis by CD73 was higher than that for AMP, and the *Vmax* value for AMP hydrolysis was higher than that for FAD hydrolysis, suggesting that CD73 binds AMP with higher affinity than FAD and more efficiently hydrolyzes AMP than FAD. In contrast, the *Km* values for FMN and AMP hydrolysis by ALP were comparable.

### Inhibition studies of ALP and CD73

Several types of inhibitors were used to characterize the hydrolytic properties of CD73 and ALP (Fig 3). FMN hydrolysis and AMP hydrolysis by ALP showed similar patterns of inhibition (Fig 3A). FAD hydrolysis and AMP hydrolysis by CD73 also showed similar patterns of inhibition (Fig 3B). However, the inhibition properties of AMP hydrolysis mediated by ALP and CD73 were different. FMN and nicotinamide mononucleotide (NMN), which are phosphorylated vitamins, and levamisole, which is an inhibitor of TNSALP (Kozlenkov *et al*, 2004), decreased the activity of ALP but not CD73. In contrast, α,β-methylene adenosine 5′-diphosphate (APCP), an inhibitor of CD73 (Bhattarai *et al*, 2015), inhibited CD73 but not ALP. GMP, a nucleotide, inhibited both CD73 and ALP, which is consistent with AMP being a common substrate of CD73 and ALP (Fig 3C). Figs 2 and 3 demonstrate that TNSALP hydrolyzes FMN, producing RF and CD73 hydrolyzes FAD, producing FMN.

**Figure 3.**
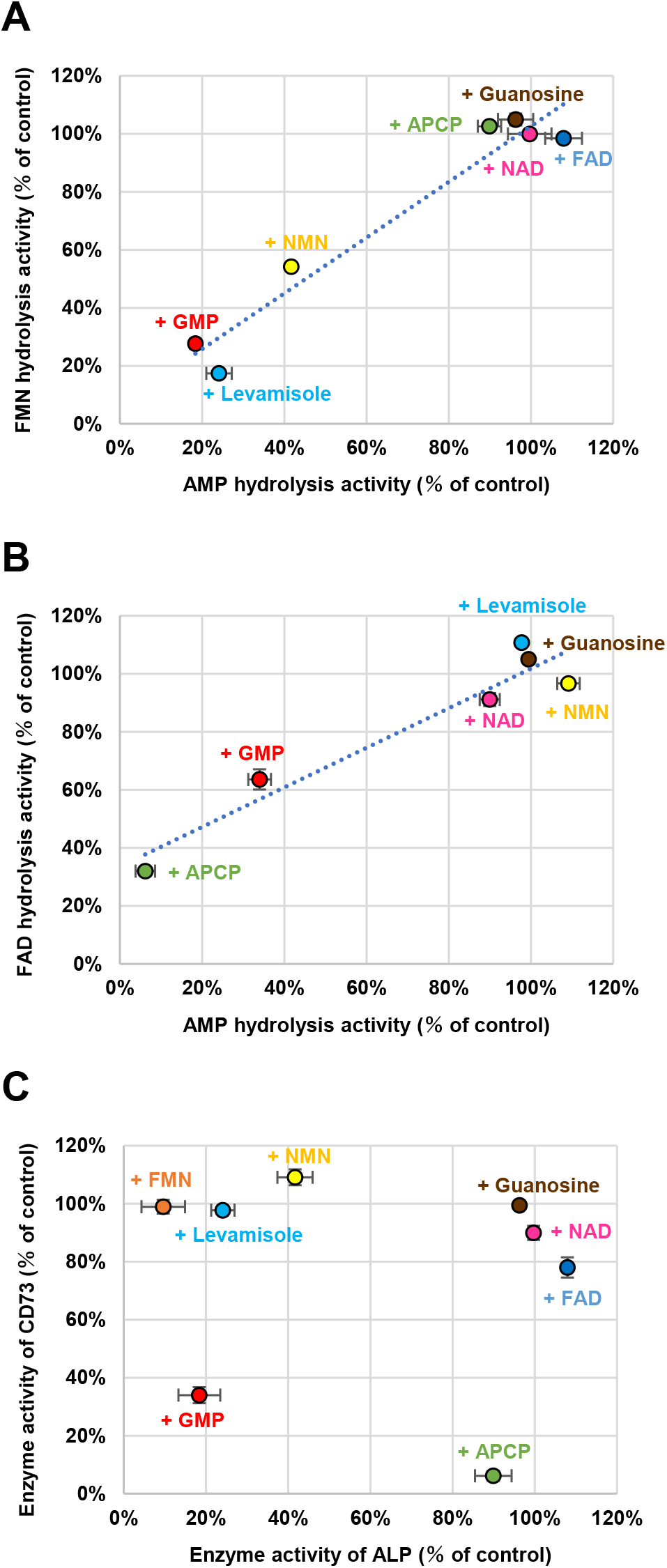
Inhibition of ALP and CD73. A. Comparison of inhibitory effect of various compounds on hydrolysis of 1 μM FMN and 1 μM AMP by ALP. FMN hydrolysis activities were measured by RF production from FMN; AMP hydrolysis activities were measured by adenosine production from AMP. The concentration of the inhibitors was 10 μM, except for α,β-methylene adenosine 5′-diphosphate (APCP; 2 μM) and levamisole (1 mM). Data represent means ± S.D. (*n* = 3). When the S.D. is smaller than the symbols, it is not shown. B. Comparison of inhibitory effects of various compounds on hydrolysis of 10 μM FAD and 4 μM AMP by CD73. FAD hydrolysis activities were measured by FMN production from FAD; AMP hydrolysis activities were measured by adenosine production from AMP. The concentration of the inhibitors was 10 μM, except for APCP (2 μM) and levamisole (1 mM). Data represent means ± S.D. (*n* = 3). When the S.D. is smaller than the symbols, it is not shown. C. Comparison of inhibitory effects of various compounds on hydrolysis of 1 and 4 μM AMP by ALP and CD73, respectively. AMP hydrolysis activities were measured by adenosine production from AMP. The concentration of the inhibitors was 10 μM, except for APCP (2 μM) and levamisole (1 mM). Data represent means ± S.D. (*n* = 3). When the S.D. is smaller than the symbols, it is not shown.

### Extracellular hydrolysis and uptake of vitamin B_2_, and vitamin B_2_-dependent PLP and PL production in GPI-deficient cells

Both CD73 and ALP are GPI-APs (Misumi *et al*, 1990; Zimmermann *et al*, 2012) and phosphatidylinositol glycan anchor biosynthesis class T (PIGT) is required for GPI-AP generation, and therefore PIGT-knockout (KO) SH-SY5Y cells (PIGT−cells) were generated using the CRISPR/Cas9 system to obtain CD73- and ALP-defective cells. SH-SY5Y is the human neuroblastoma cell line. The activities of CD73 and ALP in PIGT− cells were significantly lower than in PIGT rescued cells (PIGT+ cells) (Fig 4A, B). Surface expression levels of CD73, TNSALP, and CD59 (another GPI-AP) on PIGT− and PIGT+ cells were compared by flow cytometric analysis (Fig 4C). The surface expression of CD73, TNSALP, and CD59 was deficient on PIGT− cells.

**Figure 4.**
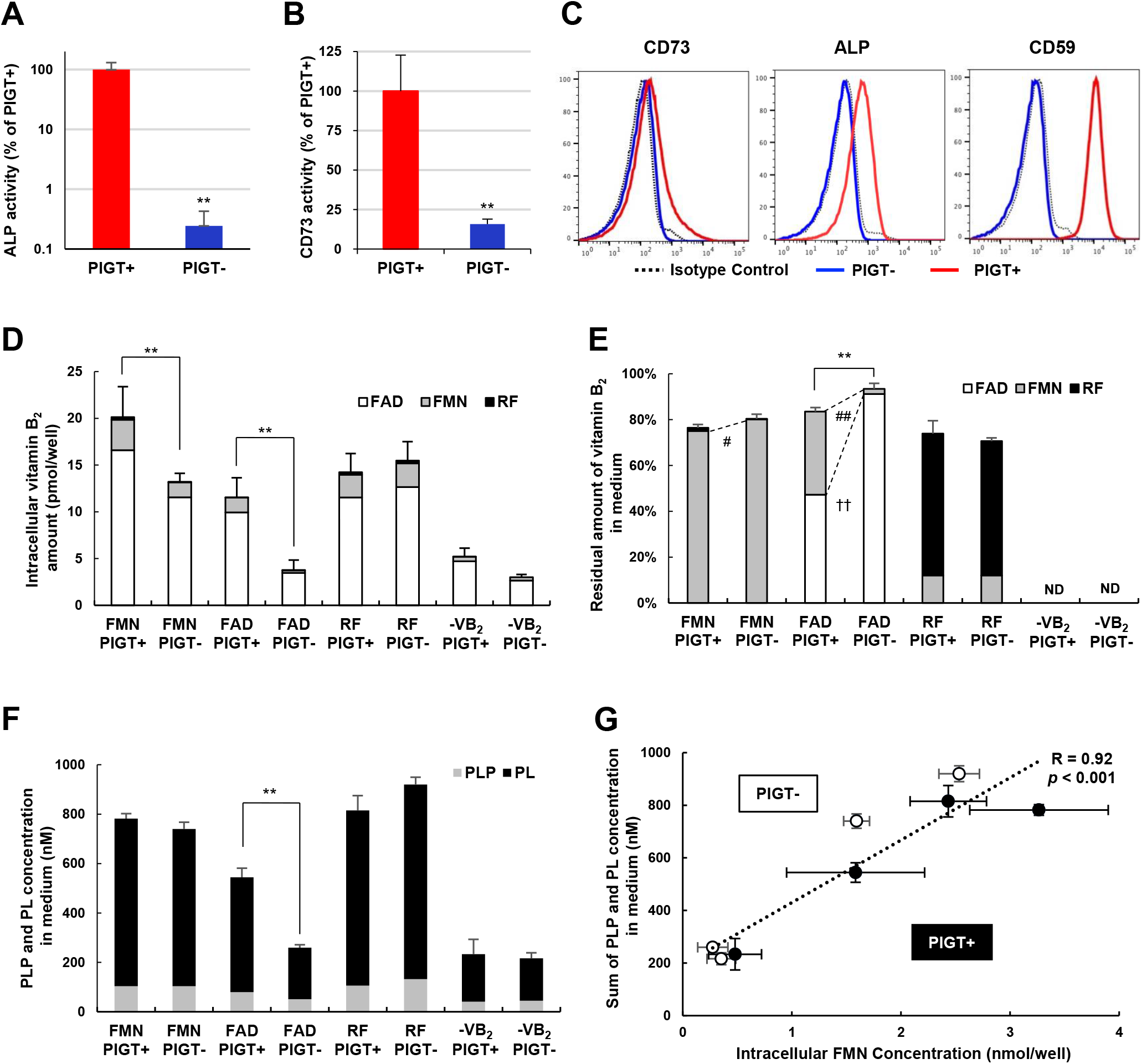
Extracellular hydrolysis and uptake of vitamin B_2_, and vitamin B_2_-dependent pyridoxal 5′-phosphate (PLP) and pyridoxal (PL) production in glycosylphosphatidylinositol (GPI)-deficient cells. A. ALP activities in phosphatidylinositol glycan anchor biosynthesis class T (PIGT)-expressing (PIGT+) or vector-transfected (PIGT−) SH-SY5Y cells. ALP activities were measured in lysates of PIGT+ and PIGT− cells. The ALP activity is expressed in terms of the amount of placental ALP (PLAP) in the kit, which was used as a positive control. Data represent means ± S.D. (*n* = 4). ** Indicates a significant difference (*p* < 0.01) compared with PIGT+ cells. B. CD73 activities in PIGT+ or PIGT− SH-SY5Y cells. CD73 activities were measured by APCP-sensitive adenosine production from AMP in lysate of PIGT+ or PIGT− SH-SY5Y cells. Data represent means ± S.D. (*n* = 3). ** Indicates a significant difference (*p* < 0.01) compared with PIGT+ cells. C. Surface expression of CD73, ALP, and CD59 on PIGT− and PIGT+ SHSY5Y cells. MFI (mean fluorescent intensity) of PIGT+ vs PIGT-; CD73, 177 vs 82; ALP, 472 vs 102; CD59, 10093 vs 77) The analysis was repeated at least three times. D. Residual amount of vitamin B_2_ in medium after 24 h of cultivation of PIGT+ or PIGT− SH-SY5Y cells in medium containing FMN, FAD, RF, or no vitamin B_2_. The residual amount of vitamin B_2_ in medium is expressed as the percentage of the total vitamin B_2_ concentration (FAD + FMN + RF) in medium incubated without cells. The data represent means for FAD (white), FMN (gray), and RF (black). Error bars represent the S.D. for total vitamin B_2_. ** Indicates a significant difference (*p* < 0.01) in the residual amount of total vitamin B_2_ in the medium. ND: Not detectable. E. Intracellular vitamin B_2_ amount after 24 h of cultivation of PIGT+ or PIGT− SH-SY5Y cells in medium containing FMN, FAD, RF, or no vitamin B_2_. The data represent means for FAD (white), FMN (gray), and RF (black). Error bars represent the S.D. for total vitamin B_2_. ** Indicates a significant difference (*p* < 0.01) in the total vitamin B_2_ concentration in cells. #, ## indicates a significant difference (*p* < 0.05, *p* < 0.01, respectively) in FMN concentration in cells. †† indicates a significant difference (*p* < 0.01) in FAD concentration in cells. ND: Not detectable. F. PLP and PL concentration in medium after 24 h of cultivation of PIGT+ or PIGT− SH-SY5Y cells in medium containing FMN, FAD, RF, or no vitamin B_2_. The data represent means for PLP (gray) and PL (black). Error bars represent the S.D. of the sum of PLP and PL. ** Indicates a significant difference (*p* < 0.01) in the sum of PLP and PL in the medium. ND: Not detectable. G. Correlation between intracellular FMN concentration and the sum of PL and PLP concentration after 24 h of cultivation of PIGT+ (closed circles) or PIGT− (open circles) SH-SY5Y cells in medium containing FMN, FAD, RF, or no vitamin B_2_. The data represent means ± S.D. (*n* = 3). Statistical analysis: Student’s t-test for (A, B); ANOVA followed by the Tukey–Kramer test for (D, E and F); Pearson correlation for (G).

PIGT− and PIGT+ cells were cultured in vitamin B_2_-depleted medium for 5 days, followed by culture for 24 h in a medium containing one of the vitamin B_2_ derivatives (FMN, FAD, or RF) or a medium without vitamin B_2_. Vitamin B_2_ concentrations in the cell and culture medium were measured by HPLC. FAD was the major form of vitamin B_2_ in the PIGT+ and PIGT− cells after cultured in medium containing RF, FMN or FAD, indicating that imported RF was intracellularly converted to FAD (Barile *et al*, 2016). The intracellular total vitamin B_2_ concentrations were significantly lower in PIGT− cells than in PIGT+ cells after cultured in FMN- or FAD-containing medium, while they were similar after cultured in RF-containing or vitamin B_2_-depleted medium (Fig 4D) (because RF can be transported into the cells by riboflavin transporters).

In the media, total vitamin B_2_ and FAD were significantly higher (91.2 ± 2.1% *vs*. 47.3 ± 0.3%, *p* < 0.01) and FMN was significantly lower (2.2 ± 0.4 % *vs*. 36.2 ± 2.0%, *p* < 0.01) for PIGT− cells cultured with medium containing FAD than for PIGT+ cells, suggesting that FAD was not efficiently hydrolyzed to FMN by PIGT− cells (Fig 4E). FMN was significantly higher (80.1 ± 2.1% *vs*. 75.0 ± 0.3%, *p* < 0.05) for PIGT− cells cultured with medium containing FMN than for PIGT+ cells, suggesting that FMN was not efficiently hydrolyzed to RF by PIGT− cells (Fig 4E). However, when cultured with medium containing RF, there was no difference in vitamin B_2_ concentration in medium between PIGT− and PIGT+ cells (Fig 4E).

These results indicate that the presence of FAD and FMN in the medium did not lead to efficient uptake of vitamin B_2_ into GPI-deficient cells; this is because FAD and FMN were inefficiently hydrolyzed to FMN and RF, respectively, because of the defective expression of CD73 and ALP; this resulted in intracellular vitamin B_2_ deficiency.

To analyze the effect of intracellular vitamin B_2_ deficiency on the activity of vitamin B_2_-dependent enzymes, concentrations of vitamin B_6_ derivatives in the medium were measured. Pyridoxine (PN, a form of vitamin B_6_) is imported into cells and phosphorylated to pyridoxine 5′-phosphate (PNP). PNP is then converted to pyridoxal 5′-phosphate (PLP) by the FMN-dependent enzyme PNP oxidase, and PLP is in turn dephosphorylated to pyridoxal (PL), PL and PLP are efficiently exported to the medium (Anderson *et al,* 1971, 1976, Da Silva *et al,* 2012). Extracellular PL and PLP should be a biomarker for intracellular vitamin B_2_ status because sum of PLP and PL are net produced amounts of metabolites by FMN-dependent PNP oxidase from PN which is contained in all precultured mediums as a vitamin B_6_ source. After cultivation of PIGT+ and PIGT− cells in the presence of PN and FAD, the combined PL and PLP concentration in the medium was significantly lower for PIGT− cells than for PIGT+ cells (Fig 4F). There was a significant positive relationship between the sum of PL and PLP concentrations in the medium and the intracellular FMN concentration (*p* < 0.001, Fig 4G). These results suggest that the amount of vitamin B_2_ imported into the cells affected the vitamin B_2_-dependent PLP and PL production.

### Effect of intracellular vitamin B_2_ deficiency on mitochondrial function

FMN and FAD act as coenzymes in the mitochondrial electron transport chain, and a decreased intracellular FAD concentration might cause mitochondrial dysfunction (Udhayabanu T et al, 2017). To compare the mitochondrial function between PIGT+ and PIGT− SHSY5Y cells, O_2_ consumption of cells was measured using a flux analyzer. After culture in vitamin B_2_-depleted medium for 5 days, cells were cultured in medium containing a vitamin B_2_ derivative (FAD, RF, or no vitamin B_2_) for 24 h and their O_2_ consumption was then measured (Fig 5). PIGT− cells showed significantly lower O_2_ consumption than PIGT+ cells when they were cultured in FAD-containing medium, whereas those cultured in RF-containing medium showed a similar level of O_2_ consumption to that in PIGT+ cells. These results suggest that the GPI-deficient cells are susceptible to mitochondrial dysfunction.

**Figure 5.**
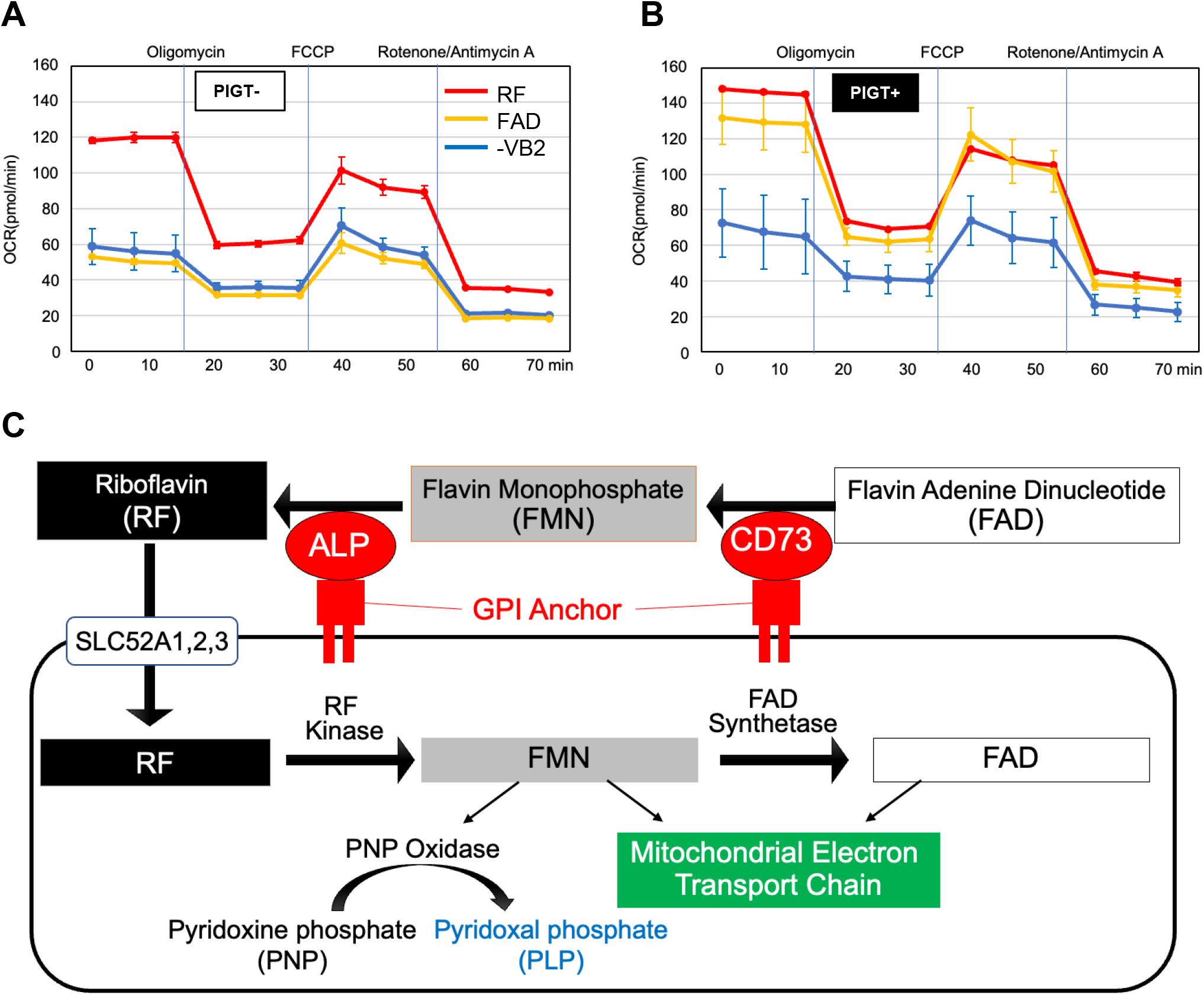
O_2_ consumption by SHST5Y PIGT− and PIGT+ cells, measured by flux analysis (A, B) and scheme of relationship between vitamin B2 metabolism and mitochondria function (C) A, B. Cells were incubated in the measuring plates with vitamin B_2_-depleted medium for 4 days. The medium was changed to the indicated conditions and the cells were further incubated for 24 h. Then, O_2_ consumption was measured. Oligomycin is an ATP synthase inhibitor; FCCP (carbonilcyanide *p*-triflouromethoxyphenylhydrazone) is an uncoupler; rotenone is a complex I inhibitor; and antimycin A is a complex III inhibitor. The data represent means ± S.D. (*n* = 2). Representative data from two independent experiments are shown. C. Scheme of hydrolysis of FAD and FMN by CD73 and alkaline phosphatase for its uptake into cells, vitamin B_6_ metabolism and mitochondria function

## Discussion

ALP catalyzes the hydrolysis of monoesters of phosphoric acid (Millán, 2006). Here, we showed that ALP hydrolyzes monophosphate vitamin B_2_, FMN, but not dinucleotide-type vitamin B_2_, FAD. Inhibition study showed that GMP and NMN inhibited ALP activity, but nicotinamide adenine dinucleotide (NAD) and FAD did not, suggesting that ALP has high affinity for compounds with a phosphomonoester moiety.

We also showed that adenosine and FMN were produced from FAD by CD73. Because AMP hydrolysis had a higher *Vmax* and lower *Km* than FAD hydrolysis by CD73, we speculate that adenosine was immediately produced from AMP after AMP and FMN production from FAD, with conversion of FAD to FMN being rate limiting. However, at the moment, we cannot completely eliminate the possibility of flavin-pyrophosphate as an intermediate.

TNSALP from human bone was used in the present study. TNSALP hydrolyzes some phosphate compounds, such as inorganic pyrophosphate, PLP (vitamin B_6_), and thiamine pyrophosphate (vitamin B_1_). Hypophosphatasia (HPP) is caused by loss-of-function mutations in TNSALP. Decreased conversion of pyrophosphate to phosphate caused dysosteogenesis (Millán & Whyte, 2016). Because cell surface hydrolysis of PLP to PL is important for vitamin B_6_ uptake into cells, decreased PLP hydrolysis activity causes vitamin B_6_ deficiency, leading to dysfunction of various vitamin B_6_-dependent enzymes such as glutamate decarboxylase which results in pyridoxine-dependent seizures (Millán & Whyte, 2016; Akiyama *et al*, 2018). In HPP, lowered levels of thiamine pyrophosphate in red blood cells were reported (Luong & Nguyen, 2005). Adenosine and GABA concentrations were lower in brain of *Akp2* KO mice than in wild-type mice; this gene encodes TNSALP in mice (Cruz *et al*, 2017). The present study showed that the phosphorylated form of vitamin B_2_, FMN, is a substrate of human TNSALP. Because hydrolysis by ALP of vitamin B_1_, B_2_, and B_6_ is required for their uptake, their uptake would be decreased in HPP, which will be a subject of future investigation.

Analysis using GPI-deficient cells, which are defective in both CD73 and ALP cell surface expression, showed that both FAD and FMN uptake activities were lower than in GPI-rescued cells, leading to intracellular deficiency of vitamin B_2_. Thus, the GPI-deficient cells showed dysfunction of the vitamin B_2_-dependent mitochondrial respiratory chain complex, as well as of PNP oxidase, and enzyme in vitamin B_6_ metabolism. GPI-deficient cells showed significantly lower PLP and PL production and O_2_ consumption when they were incubated with FAD than PIGT+ cells, whereas those incubated with RF showed a similar level to that in PIGT+ cells (Figs 4F and 5A, B). In addition, a significant positive relationship was found between the concentration of intracellular FMN and the sum of PLP and PL production (Fig 4G). These results again suggest that cell surface hydrolysis of FAD to RF by the CD73 and ALP contributed to vitamin B_2_-dependent functions (Fig 5C).

IGD is caused by mutations in genes involved in the biosynthesis or modification of GPI-APs. Major symptoms of patients with IGD are intellectual disability, developmental delay, and seizures. Because both ALP and CD73 are GPI-APs, expression of these proteins is decreased in some patients with IGD. Here, we demonstrated that GPI-deficient cells showed decreased intracellular vitamin B_2_ levels when cultured with FAD, a major form of vitamin B_2_ in blood, which led to mitochondrial dysfunction. This is consistent with reports that some severe IGD cases show mitochondrial dysfunction (Tarailo-Graovac *et al*, 2015). CD73 expression is decreased in phosphatidylinositol glycan anchor biosynthesis class G (PIGG) knockout cells (Ishida *et al*, 2022) and some cases with null mutation of PIGG also showed decreased expression of CD73 and mitochondrial dysfunction (Tremblay-Laganière *et al*, 2021), suggesting that CD73 expression is important for uptake of vitamin B_2_ in PIGG deficiency. Similar to HPP, some patients with IGD show decreased vitamin B_6_ uptake and suffer from pyridoxine-dependent seizures (Kuki *et al*, 2013). High-dose non-phosphorylated vitamin B_6_ (i.e., pyridoxine) treatment was effective in treatment of seizures in more than half of patients with IGD (Tanigawa *et al*, 2021). Two types of non-phosphorylated vitamin B_6_ are imported into cells and converted to PLP in the cell. One of them, PL, is contained in food from animal sources. Pyridoxine and its glycoside are contained in food from plant sources (Gregory & Ink, 1987); pyridoxine is intracellularly converted to PNP, and PNP is converted to PLP by FMN-dependent PNP oxidase (Anderson, 1971, 1976, Da Silva *et al*, 2012). In addition to decreased uptake of vitamin B_6_, patients with IGD would show decreased intracellular conversion of PNP to PLP due to dysfunction of this FMN-dependent enzyme caused by the decreased vitamin B_2_ uptake. Therefore, high-dose pyridoxine and riboflavin treatment might be effective for patients with IGD. However, further study is required concerning vitamin B_6_ and B_2_ metabolism in patients with IGD.

Adenosine, produced by CD73 from AMP, is an immune inhibitory molecule through its receptor expressed on immune cells (Stagg *et al*, 2010; Allard *et al*, 2017). Some tumors show upregulation of CD73, and adenosine promotes both migration and proliferation (Wang *et al*, 2011). Therefore, CD73 is a target for immunotherapy for cancer. Antibody against CD73 has been used in a phase 1 clinical study (Allard *et al*, 2017). Here, we show the importance of CD73 for vitamin B_2_ uptake. In cancer therapy the effect of antibody against CD73 on vitamin B_2_ metabolism should be studied in future study.

## Materials and Methods

### Materials

Human bone ALP was purchased from CALZYME (San Luis Obispo, CA). Human CD73 His-tag was purchased from BPS Bioscience (San Diego, CA). All other reagents were of analytical grade.

### Measurement of hydrolysis of FAD

For measurement of hydrolysis of FAD, FAD and human bone ALP or human CD73 were incubated in reaction buffer containing 50 mM Tris-HCl (pH 7.4), 5 mM MgCl_2_, and 0.002% bovine serum albumin, after separately preincubation of FAD and the enzyme at 37°C for 3 min. At an appropriate time, the reaction was stopped by addition of ice-cold perchloric acid and the mixture was centrifuged (9,000 × *g* for 5 min). The supernatant was transferred to a fresh tube, neutralized by KOH, and recentrifuged. The supernatant was filtered through a 0.45-μm membrane and simultaneously analyzed for FMN, RF, AMP, and adenosine concentrations by HPLC. The HPLC conditions were optimized from a previously reported method (Akimoto *et al*, 2006). The HPLC apparatus consisted of a Shimadzu LC-10ADvp system equipped with an RF-10Axl spectrofluorometer and SPD-10Avp UV-VIS detector (Shimadzu, Kyoto, Japan).

Chromatographic separation was performed on an InertSustain AQ-C18 column (150 × 4.6 mm, i.d. 5 μm; GL Sciences, Tokyo, Japan) using a gradient elution mode at a flow-rate of 1.0 mL/min. The column temperature was 25°C. Mobile phase A was 10 mM potassium phosphate containing 5 mM ethylenediaminetetraacetic acid-disodium salt, adjusted to pH 6.0; mobile phase B was methanol. We used a linear gradient from 8% to 25% mobile phase B from 3 min to 6 min, followed by holding at 25% B until 25 min after injection. Fluorescence measurements were made with excitation at 440 nm and emission at 560 nm to determine FMN and riboflavin concentrations. Absorbance measurements were made at 260 nm for the determination of AMP and adenosine. Hydrolysis activity was calculated as observed production *minus* non-enzymatic production, which was determined in negative controls from which enzyme was absent otherwise using the same procedure as described above.

### Measurement of hydrolysis of FMN

For measurement of hydrolysis of FMN, FMN and human bone ALP or human CD73 were incubated in reaction buffer containing 50 mM Tris-HCl (pH 7.4) and 5 mM MgCl_2_ at 37°C for 3 min (Coburn *et al*, 1998). The reaction procedures were the same as for analysis of FAD hydrolysis. The produced RF concentration was determined using an isocratic HPLC method with 75% buffer A and 25% buffer B (other conditions the same as for analysis of FAD hydrolysis).

### Measurement of hydrolysis of AMP

For measurement of hydrolysis of AMP, AMP and human bone ALP or human CD73 were incubated in reaction buffer containing 50 mM Tris-HCl (pH 7.4) and 5 mM MgCl_2_ at 37°C for 3 min (Coburn *et al*, 1998). The reaction procedures and HPLC conditions were the same as those for analysis of FAD hydrolysis.

### Generation of GPI-deficient cells

*PIGT*-KO cells were generated from SHSY5Y cells (a human neuroblastoma-derived cell line) using the CRIPR/Cas9 system. Plasmid pX330 for expression of human-codon-optimized *Streptococcus pyogenes* (Sp) Cas9 and chimeric guide RNA were obtained from Addgene (Cambridge, MA). The seed sequence for the SpCas9 target site in the target gene was tcggtgcagaccacctcccg<u>cgg (underline; PAM sequence)</u>. SHSY5Y cells were transfected with pX330 containing the gRNA of the target site using Lipofectamine 2000 (Invitrogen, Carlsbad, CA). KO clones were obtained by limiting dilution and KO was confirmed by sequencing the target sites in the genomic DNA and by flow cytometric analysis of CD59 expression [staining with mouse anti-hCD59 antibody (5H8) followed by phycoerythrin-conjugated anti-mouse IgG (Biolegend, San Diego, CA)]. *PIGT*-KO clone #3 was rescued by transfecting human PIGT-expressing vector pME-puro-3HA-hPIGT. After puromycin selection, the restored population was sorted to obtain PIGT+ cells. *PIGT*-KO clone #3 was transfected with an empty vector, pME-puro, and the puromycin-resistant population was selected as PIGT− cells. These cells were cultured in Dulbecco’s Modified Eagle’s Medium supplemented with 10% fetal bovine serum containing 3 μg/mL puromycin. For flow cytometric analysis, cells were stained with phycoerythrin-conjugated anti-human TNAP (B4-78, isotype; mouse IgG1, Santa Cruz Biotechnology, Dallas, TX), or anti-human CD73 (AD2, isotype; mouse IgG1, Biolegend) and anti-human CD59 antibody (isotype; mouse IgG1, clone 5H8) followed by phycoerythrin-conjugated goat anti-mouse IgG. Stained cells were analyzed using a MACS QuantVYB analyzer (Miltenyi Biotech, <u>Bergisch Gladbach</u>, Germany).

### Measurement of ALP and CD73 activities in SHSY5Y cell lines

ALP activities were measured using a Great EscAPe™ SEAP kit (Takara Bio Inc., Shiga, Japan) in lysates of PIGT+ and PIGT− cells. The ALP activity is expressed in terms of the amount of PLAP in the kit, which was used as a positive control. CD73 activities were measured by adenosine formation from AMP in lysate from PIGT+ and PIGT− cells. The quantitative method for adenosine using HPLC was described above in the section “Measurement of hydrolysis of AMP.” The APCP sensitivity of CD73 activity was calculated by subtracting the activity in the presence of 4 μM APCP from the total activity in the absence of APCP.

### Extracellular hydrolysis and uptake of vitamin B_2_, and vitamin B_2_-dependent PLP and PL production in GPI-deficient cells

PIGT− and PIGT+ cells were cultured in vitamin B_2_-deficient medium for 5 days, followed by culture in medium containing a vitamin B_2_ derivative (0.2 μM FMN, FAD, RF, or no vitamin B_2_) for 24 h. Vitamin B_2_ and B_6_ concentrations were measured in medium and cells by HPLC. The HPLC conditions for measurement of FAD, FMN, and RF were the same as described above for measurement of FMN hydrolysis. For the measurement of PL and PLP, a previously reported HPLC method with a fluorescence detector was used after precolumn derivatization with semicarbazide (Kobayashi *et al*, 2015).

### Measurement of mitochondrial function

Cellular respiration (oxygen consumption rate, OCR) was assessed using an XFp Extracellular Flux Analyzer (Seahorse Bioscience, Billerica, MA). Cells were cultured in vitamin B_2_-depleted medium for 4 days. Then, 10^4^ cells per well were incubated in poly-L-lysine (Sigma-Aldrich Inc. St. Louis, MO)-coated wells with vitamin B_2_-depleted medium, or with that supplemented by riboflavin, FAD or FMN (0.2 µM), for 24 h. The XF Cell Mito Stress Test (Seahorse Bioscience Inc.) was used to measure the key parameters of mitochondrial respiration using specific mitochondrial inhibitors and uncouplers: oligomycin (1 μM), carbonilcyanide *p*-triflouromethoxyphenylhydrazone (FCCP; 2 μM), and a mixture of rotenone/antimycin A (both 0.5 μM) were injected sequentially following the manufacturer’s instructions. Before drug addition, basal OCR was measured. Oligomycin was injected to inhibit ATP synthase (complex V), and the OCR was recorded. To determine the maximal respiration, the uncoupler FCCP was injected. Finally, a mixture of rotenone/antimycin A was injected to inhibit the flux of electrons through complexes I and III and to enable calculation of the spare respiratory capacity.

### Statistical analysis

Data are expressed as means ± S.D. Statistically significant differences were determined using one-way analysis of variance followed by Tukey’s *post-hoc* test or Student’s *t*-test, with *p* < 0.05 or 0.01 as the criterion. Pearson’s correlation analysis was performed to analyze correlations. Kinetic analyses were performed using Sigma Plot (Systat Software Inc., San Jose, CA).

## List of abbreviations

ALP: alkaline phosphatase
APCP: α,β-methylene adenosine 5′-diphosphate
FAD: flavin adenine dinucleotide
FCCP: carbonilcyanide p-triflouromethoxyphenylhydrazone
FMN: flavin mononucleotide
GPI: glycosylphosphatidylinositol
GPI-AP: GPI-anchored protein
HPLC: high-performance liquid chromatography
HPP: hypophosphatasia
IGD: inherited GPI deficiency
NAD: nicotinamide adenine dinucleotide
NMN: nicotinamide mononucleotide
PIGG: phosphatidylinositol glycan anchor biosynthesis class G
PIGT: phosphatidylinositol glycan anchor biosynthesis class T
PL: pyridoxal
PLAP: placental ALP
PLP: pyridoxal 5′-phosphate
PN: pyridoxine
PNP: pyridoxine 5′-phosphate
RF: riboflavin
TNSALP: tissue nonspecific ALP

## Acknowledgements

We thank Keiko Kinoshita, Saori Umeshita, and Kae Imanishi (Osaka University) for technical help. This work was supported by JSPS and MEXT KAKENHI grants (JP21H02415 and JP17H06422 to T. Kinoshita), a grant from Mizutani Foundation for Glycoscience, KOSE Cosmetology Research Foundation, Ministry of Health, Labour and Welfare (20FC1025), and a grant from the Practical Research Project for Rare/Intractable Diseases from the Japan Agency for Medical Research and Development (AMED) (21ek0109418h0003) to Y. Murakami. We thank Edanz (https://jp.edanz.com/ac) for editing a draft of this manuscript.

## Author contributions

N.S., D.K., T.K., N.I., K.W., and Y.M. conceived and designed the experiments. N.S., D.K., A.I., I.H., R.F., K.W., and Y.K. performed enzyme activity assays and analyzed the data. T.K. and Y.M. generated cell lines, prepared cells and performed flow cytometric analysis. N.S., D.K., R.F., N.H., and Y.M. performed analysis using cell lines. N.S., D.K., T.K., N.I., N.H., and Y.K. wrote the manuscript. All authors reviewed the manuscript.

